# Photosymbiosis reduces the environmental stress response under a heat challenge in a facultatively symbiotic coral

**DOI:** 10.1101/2023.11.13.566890

**Authors:** DM Wuitchik, HE Aichelman, KF Atherton, CM Brown, X Chen, L DiRoberts, GE Pelose, CA Tramonte, SW Davies

## Abstract

The symbiosis between corals of the order Scleractinia and dinoflagellates of the family Symbiodiniaceae is sensitive to environmental stress. The oxidative bleaching hypothesis posits that extreme temperatures lead to accumulation of photobiont-derived reactive oxygen species ROS, which exacerbates the coral environmental stress response (ESR). To understand how photosymbiosis modulates coral ESRs, these responses must be explored in hosts in and out of symbiosis. We leveraged the facultatively symbiotic coral *Astrangia poculata*, which offers an opportunity to uncouple the ESR across its two symbiotic states (symbiotic, aposymbiotic). Colonies of both symbiotic states were exposed to three temperature treatments for 15 days: i) control (static 18°C), ii) heat challenge (increasing from 18 to 32°C), and iii) cold challenge (decreasing from 18 to 6°C) after which host gene expression was profiled. Cold challenged corals elicited widespread differential expression, however, there were no differences between symbiotic states. In contrast, symbiotic colonies exhibited greater gene expression plasticity under heat challenge, including enrichment of cell cycle pathways involved in controlling photobiont growth. Counter to the oxidative bleaching hypothesis, this plasticity did not include signatures of stress, and rather a dampened ESR under heat challenge was observed, suggesting that photobionts reduce the host’s ESR under elevated temperatures in *A. poculata*.

## Introduction

The photosymbiosis between coral hosts and endosymbiotic algae in the family Symbiodiniaceae forms the backbone of entire coral reef ecosystems. This symbiosis is particularly important as tropical reef-building corals are found in nutrient poor waters, which are often carbon and nitrogen limited [1]. Much of the organic carbon required by coral hosts comes from translocation of photosynthetically derived sources from the symbiotic algae (photobiont) [2] and hosts can actively promote photosynthesis through acidification of the symbiosome where photobionts reside [3]. Nitrogen limitation is overcome in part through interactions between the host and photobiont that conserve and recycle nitrogen between partners [4]. Coral-algal photosymbiosis is therefore vital to a healthy coral reef ecosystem; however, this relationship is vulnerable to rising seawater temperatures associated with anthropogenic climate change, and warmer temperatures lead to the breakdown of photosymbiosis (*i.e.,* dysbiosis) in a process termed coral bleaching [5,6]. Bleaching can lead to coral mortality as hosts are deprived of symbiont-derived carbon sugars [7], and bleaching events have led to dramatic declines in coral reefs globally [8], negatively impacting coastal communities [9,10]. While rising temperatures are the most immediate threat to coral reefs, cold water extremes also cause coral bleaching and pose significant thermal challenges to coral species that are rarely investigated alongside rising temperatures [11–13]. As coral bleaching episodes become more frequent and severe as climate change accelerates [14], understanding the mechanisms underlying this dysbiosis has become increasingly important.

Coral bleaching research has largely focused on corals that exhibit obligate symbiotic relationships (for review, see [15]). These works have highlighted the importance of heat-shock proteins [16,17], antioxidant pathways [18], and immunity [19,20] in the coral environmental stress response (ESR). A recent meta-analysis of transcriptomic stress responses in *Acropora* corals compared gene expression profiles from 14 distinct experiments and found that high-intensity stressors elicited stereotyped ESRs, with high stress ESRs leading to a ‘Type A’ response whereas corals experiencing less severe stressors show the opposite ESR pattern (‘Type B’ response) [21]. These characterisations have been a useful tool in ascertaining whether experimental treatments elicit similar ESR across gene expression studies in corals [22]. While we now have a broad understanding of coral ESRs in obligate symbiotic hosts, these corals cannot survive without their photobionts for extended time periods due to nutritional constraints [2]. Therefore, disentangling the effects of temperature from those associated with nutritional stress when photosymbiosis is lost remains a challenge, leaving critical gaps in our understanding of how photosymbiosis modulates coral ERSs.

Photosymbiosis may alter the coral host ESR depending on the context of the stressor. The oxidative bleaching hypothesis posits that the photosynthetic machinery of the photobiont malfunctions under increasing temperatures, leading to an accumulation of photobiont-derived reactive oxygen species (ROS), ultimately exacerbating the coral ESR [23]. This pattern of photoinhibition followed by ROS accumulation has been suggested for cold thermal stress as well [24]. However, ROS accumulation has been found in the host preceding photosynthetic dysfunction [25], and hosts have been shown to exhibit stronger stress-related gene expression responses than their photobionts [19,26]. Furthermore, photobionts expelled during bleaching are photosynthetically competent [27], which challenges the notion that photo-oxidative stress is the weak link leading to dysbiosis. Alternatively, photobionts may provide critical energy reserves necessary for the host’s ESR [28]. For example, production of heat shock proteins that aid in repairing cell damage is energetically costly [29] and additional energy reserves may mitigate bleaching. Indeed, heterotrophy can reduce the probability of dysbiosis [30,31]. Ultimately, to understand if photosymbiosis modulates the ESR and supports the oxidative bleaching hypothesis (Table 1), these responses must be explored in coral hosts in and out of symbiosis.

**Table 1.** Predictions based on the oxidative bleaching hypothesis in how photosymbiosis impacts the coral environmental stress response (ESR) during thermal challenges.

| Oxidative Bleaching Hypothesis | **H0a**<br>No effect of photosymbiosis | **H0b**<br>Photosymbiosis decreases ESR | **H1**<br>Photobiont derived ROS increases ESR |
| --- | --- | --- | --- |
| Differential expression between symbiotic states under thermal challenge | No | Yes | Yes |
| Key expression signatures | NA | No change in oxidative stress under thermal challenge | Increase in oxidative stress symbiotic coral |
| ESR (Dixon *et al.* 2020) | No association | Type B | Type A |

Coral-algal photosymbioses exist along a continuum, with some coral species being completely heterotrophic (aposymbiotic) and others being fully reliant on autotrophy of their photobionts (symbiotic) [32]. Facultatively symbiotic corals exist across this continuum, offering the opportunity to disentangle host responses in and out of symbiosis by leveraging the aposymbiotic and symbiotic phenotypes, which occur naturally on subtropical and temperate reefs [33–39]. These phenotypes correspond to photobiont densities, with symbiotic colonies having higher photobiont densities than aposymbiotic colonies (aposymbiotic colonies < 1 × 10^5^/cm^2^; [40]). Despite aposymbiotic colonies having background populations of photobionts, these levels have a *de minimis* physiological effect on coral hosts (see [41]). Leveraging these symbiotic states in facultative corals has been useful in exploring key molecular pathways of photosymbiosis maintenance, including nitrogen cycling [37], symbiont density regulation, and immunity [42]. The facultatively symbiotic coral, *Astrangia poculata,* has emerged as a model system for symbiosis research [37,43,44] and previous gene expression work from our group found that aposymbiotic *A. poculata* exhibit classic ESRs consistent with those observed in tropical reef-building corals [22].

Here, we build on this work by leveraging aposymbiotic and symbiotic colonies of *A. poculata* to explore how symbiosis modulates the host’s ESR with the goal of testing the oxidative bleaching hypothesis. Under this hypothesis, we would expect an elevated ESR in symbiotic corals due to the accumulation of photobiont derived ROS, and a more muted ESR in aposymbiotic hosts (Table 1). We exposed aposymbiotic and symbiotic *A. poculata* to both cold and heat challenges and sampled for gene expression to examine molecular snapshots of how symbiosis modulates the host’s ESR. To gain deeper insights into these responses, we compare these ESR profiles with those observed in tropical reef-building corals. Together, these data provide insights into the molecular repertoires a facultative coral host in and out of symbiosis engages to withstand thermal challenges.

## Materials and Methods

### Astrangia poculata thermal challenge experiment

*Astrangia poculata* colonies (N=20) were collected from Woods Hole, MA, USA (N41° 31.51, W70°40.49; Figure 1) and shipped overnight to Boston University. Each colony was cut into three fragments, attached to petri dishes using cyanoacrylate glue, and maintained at 18°C for several months of recovery. Colonies were classified by phenotype as either aposymbiotic (white) or symbiotic (brown) and then randomly assigned to one of three treatments (Table 1). Each treatment consisted of three replicate experimental tanks with temperatures maintained by Aqualogic Digital Temperature Controllers connected with aquarium heaters and chillers. A 12:12 h light:dark cycle was maintained with an intensity of 50 µmol m^-2^ s^-1^ and fragment positions were systematically rotated throughout to avoid potential differences in light exposure and water flow. Temperature and salinity were monitored daily using a YSI pro30 multiprobe and these values were confirmed with a glass thermometer and refractometer. The experiment was run for 15 days and control tanks were maintained at 18°C (salinity 35.3 +/- 0.6 ppt) for the duration of the experiment. Tanks in the cold challenge were cooled from 18°C by approximately 1°C per day until a final temperature of 4°C was achieved (salinity 34.5 +/- 1.5 ppt), and tanks in the heat challenge were heated from 18°C by approximately 1°C per day until a final temperature of 30°C was reached (salinity 35.0 +/- 1.1 ppt) (Figure 1B,D). Hourly temperature data between 2014-2021 were obtained from the National Oceanic and Atmospheric Administration (NOAA) weather buoy number BZBM3 and plotted with experimental challenge temperatures to compare treatments relative to collection site temperatures (Figure 1C).

**Figure 1.**
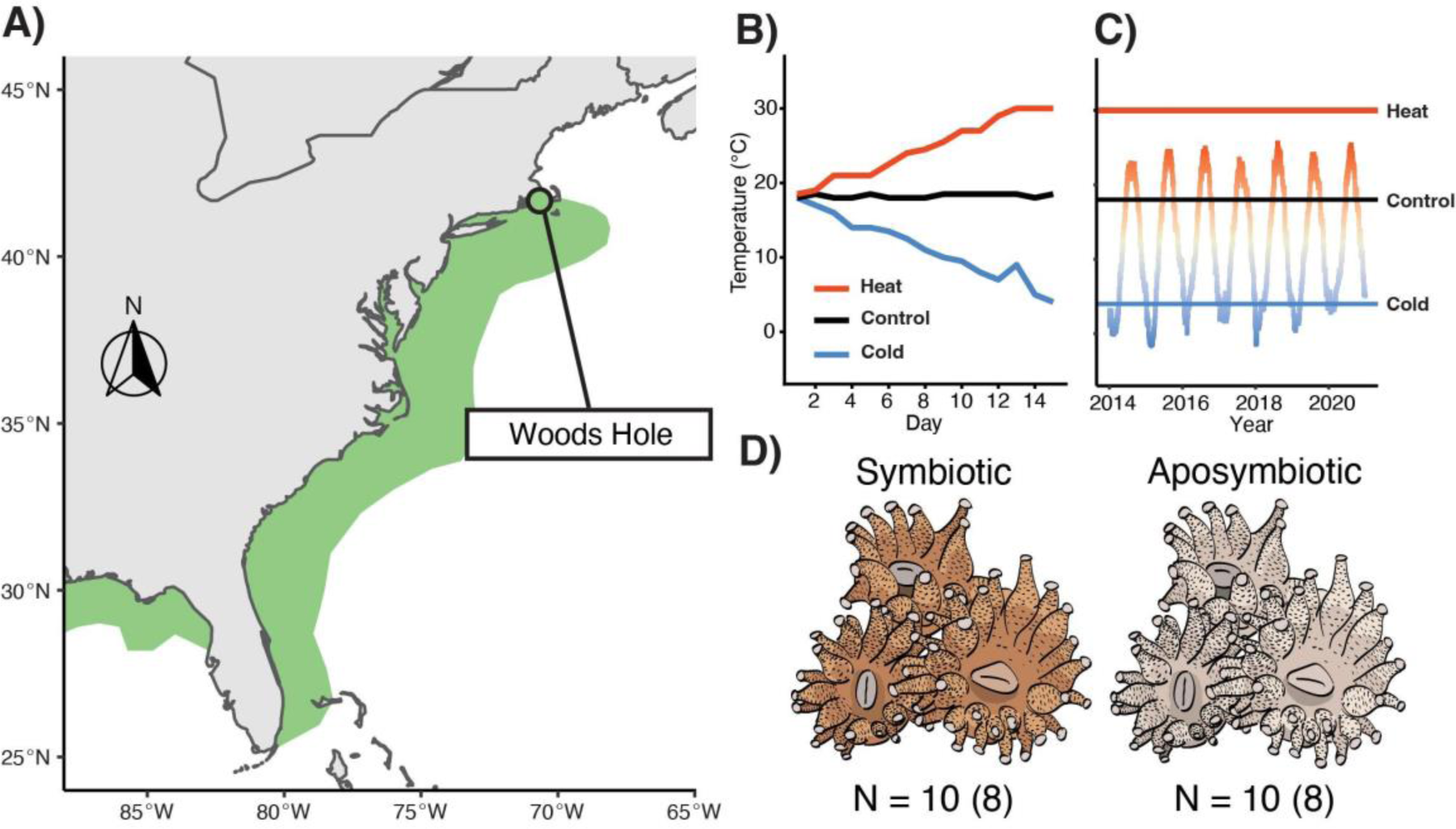
*Astrangia poculata* thermal challenge experimental design. A) Map of the eastern seaboard of the United States with documented *Astrangia poculata* distributions in green (distributions based on Thornhill et al., 2008). B) Temperature data for three experimental treatments throughout a 15 day thermal challenge. C) Mean hourly temperature profiles at Woods Hole, MA with final temperature reached in each challenge treatment overlayed (Black: Control, Red: Heat, Blue: Cold). Colour gradient shifts from red (maximum temperature) to grey (mean temperature) to blue (minimum temperature). Seasonal temperature data were obtained from the National Oceanic and Atmospheric Administration (NOAA) weather buoy number BZBM3. D) Symbiosis phenotypes of *A. poculata* with sample sizes used for behavioural assay (gene expression analyses in parentheses).

### Coral behavioural response to food stimuli

To determine behavioural responses to temperature challenge and symbiotic state, coral polyp behaviours in response to food stimulus were quantified following previous studies [22,33]. In brief, polyp activity was scored daily on a scale of 1 to 5 based on the percentage of active polyps within a fragment (1 = 0%, 2 = 25%, 3 = 50%, 4 = 75%, 5 = 100%) 30 minutes after freeze-dried copepods (Argent Cyclop-Eeze) were suspended in seawater. The same observer conducted each assay to limit observer biases. A cumulative link mixed model for ordinal-scale observations using the R package *ordinal* [45] was generated with genotype and experimental tanks as random effects, and temperature treatment, symbiotic state and experimental day as fixed effects.

### Tag Seq library preparation and sequencing

Upon reaching maximum differences between thermal challenge and control treatments (Day 15), polyps from replicate colonies (Nsymbiotic = 8, Naposymbiotic = 8) were sampled from each treatment (Ntotal samples = 48) for gene expression profiling. Several polyps were removed from each fragment using sterilized bone cutters, immediately preserved in 200 proof ethanol, and stored at –80°C. Total RNA was extracted using an RNAqueous kit (Ambion by LifeTechnologies) following manufacturer’s recommendations. An additional step was implemented using 0.5 mm glass beads (BioSpec), which were added to the lysis buffer and samples were homogenized using a bead beater for 1 min. RNA quantity was determined using a DeNovix DS11+ spectrophotometer and integrity was assessed by visualising ribosomal RNA bands on 1% agarose gels. Trace DNA contamination was removed using a DNase 1 (Ambion) digestion at 37°C for 45 min. Libraries were created from 1500 ng of total RNA following Meyer et al., [46] and adapted for Illumina Hi-Seq sequencing [47]. In brief, RNA was heat sheared and transcribed into first strand cDNA using a template switching oligo and SMARTScribe reverse transcriptase (Clontech). cDNA was PCR amplified, individual libraries were normalized, and Illumina barcodes were incorporated using a secondary PCR. Samples were pooled and size-selected prior to sequencing on Illumina Hiseq 2500 (single-end 50 base pairs) at Tufts University Core Facility.

### Gene expression analyses

References for *A. poculata* [48] and its homologous photobiont *Breviolum psygmophilum* [42] were concatenated to form a holobiont reference. Raw sequence files were trimmed to remove adapters and poly-A tails using the *fastx-Toolkit* (v 0.0.14, Gordon & Hannon, (2010) fastx-Toolkit. http://hannonlab.cshl.edu/fastx_toolkit.) and sequences that were <20 bp in length were removed. Sequences with >90% of bases having a quality score >20 were retained, PCR duplicates were removed, and resulting quality-filtered reads were mapped to the holobiont reference using *bowtie2* v2.4.2 [49]. Samples maintained in analyses had an average of 703,688 (SD = 298,412) mapped reads and five individuals were removed due to low read depth (< 100,000/sample, Table S1). Of the remaining samples, mapping efficiencies ranged from 37%-83% with an average mapping efficiency of 69% (SD = 9.4%) (Table S2). Reads that were assigned to the photobiont (1042-41035 total reads; 0.1-10.0 %) were then discarded and only host reads were used for subsequent analyses.

The presence of clones in the dataset was checked by identifying single nucleotide polymorphisms (SNPs) across samples. Reads from each sample were mapped to the host genome using *bowtie* v2.4.2 [49], which produced sam alignment files that were converted to bam files using *sortConvert* in *samtools* [50]. An identity-by-state matrix was then generated using *ANGSD* v.0935 [51] with loci filtered to include those present in 78% of individuals, having a minimum quality score of 25 and a mapping score of 20. Further parameters in *ANGSD* were set so that a strand bias p-value > 1 x10-5, minimum minor allele frequency > 0.05, p-value > 1 x 10-5, excluding all triallelic sites as well as reads with multiple best hits. A dendrogram was created using *hclust* v 3.6.2 and samples that were separated by a height of less than 0.15 were classified as clonal given that this height clustered replicate fragments of the same genet (Figure S1). No clones were observed; however, one sample (AP4) was removed from further analyses due to its high divergence from all other samples in the dataset, which was strong evidence that it was an outlier (Figure S2)

Further outlier examination was conducted using *arrayQualityMetrics* in *DESeq v1.39* [52]. One sample was flagged as an outlier and another was identified as having a high likelihood of being mislabelled (Figure S2, AC1). Both samples were removed from subsequent analyses (Table S2). To determine differentially expressed genes (DEGs), *DESeq2* v1.30 [53] was used with a correction for multiple testing done using the Benjamini and Hochberg method (FDR < 0.05) [54]. To test how symbiotic state modulated the gene expression response, we conducted pairwise contrasts between either heat or cold challenge with control corals for each symbiotic state separately. Lists of these DEGs isolated from each contrast were compared between symbiotic states using a Venn diagram. This process generated a list of unique and intersecting DEGs, which were used to perform a series of gene ontology (GO) enrichment analyses using Fisher’s exact tests within the GO_MWU R package [55].

Global gene expression patterns were assessed by performing a variance stabilizing transformation (vst) followed by a principal component analysis (PCA). This PCA was then tested for differences between treatment levels using a permutational multivariate analysis of variance with the *adonis* function in *vegan v2.5.4* [56]. In addition to the PCA, these findings were further evaluated by r-log transforming gene expression data followed by a discriminant function analysis (DAPC). Both the PCA and DAPC were given a gene expression plasticity score using a custom function [57] based on distance between samples in the first two PC axes relative to the mean of all control samples. The effects of symbiotic state and treatment on gene expression plasticity on both the PCA and DAPC were tested by first checking for assumptions of normality and equal variance followed by an ANOVA and Tukey’s honest significant differences post hoc tests.

### Colour Analysis

Because *A. poculata* exists along a symbiosis continuum, we assessed the strength of our categorical symbiotic state assignments, which were initially judged visually using brown (symbiotic) or white (aposymbiotic) phenotypes. Photos of each coral fragment were taken at the beginning of the experiment using a Coral Watch Coral Health reference card as a standard for light exposure [26]. All images were white balanced using Adobe Photoshop and then ten points on the coral were randomly selected using ImageJ. The red channel intensity was calculated from these points using custom MATLAB script [58] as a proxy for photobiont density. As an additional proxy, we explored the relative abundance of sequence reads that mapped to the photobiont transcriptome relative to reads that mapped to the host genome. To determine whether these photobiont reads corresponded with symbiotic state, we first transformed the percentage of photobiont reads to meet assumptions of normality using a square root transformation. Next, we fit a linear model (formula: sqrt(Percent mapped to photobiont) ∼ Treatment + Symbiotic State + Treatment:Symbiotic State) to test whether symbiotic state and treatment could predict the mapped photobiont counts. Lastly, we performed an additional linear model between percentage of square root transformed photobiont reads and red channel intensity to test whether the percentage of photobiont reads served as a proxy for coral colour.

### Evaluating the coral environmental stress response (ESR)

We first performed GO enrichment analysis using a Mann-Whitney U test (GO-MWU) based on rankings of signed log p-values [59] for the heat and cold thermal challenges separately. In these analyses, we set parameters to filter GO categories if they contained more than 10% of the total number of genes, contain at least 10 genes to be considered, and the cluster cut height set to 0.01 to suppress merging of GO terms. This provides delta-ranks of each GO term, which quantifies the tendency of associated genes as being up- or downregulated in challenge samples vs controls. To evaluate the oxidative bleaching hypothesis, we looked for signals of a response to ROS by isolating the children GO terms under the parent term *oxidative stress* (GO:0006979) using *GOfuncR v 1.10.0* [60]. This broadly captures GO terms associated with response to ROS and the average delta ranks were compared between treatments and symbiotic states using an ANOVA with fixed effects of treatment and symbiotic state.

To characterise how the environmental stress response (ESR) differs across symbiotic states in each of the thermal challenges, we compared GO enrichment values from our data with results of a meta-analysis isolating the ESR from the genus *Acropora* [21]. While not a formal statistical analysis due to GO being non-independent from each other (overlapping gene sets), this presents a qualitative way to compare functional similarity between experiments. Therefore, a positive relationship would indicate that the ESR is largely consistent with the Type “A” ESR and a negative relationship would suggest a Type “B” ESR. This comparison determines whether our thermal challenges elicits responses consistent with those observed in previous work conducted in tropical corals.

## Results

### Confirmation of symbiotic state assignment

Aposymbiotic colonies had significantly lower proportions of counts mapping to the *B. psygmophilum* transcriptome than symbiotic colonies (beta = -0.97, 95% CI [-1.46, -0.48], t(33) = -3.87, p < .001). We then fit a linear model to predict the proportion of counts mapping to the *B. psygmophilum* transcriptome with the red intensity calculated from the image colour analysis. This resulted in a significant negative relationship explaining 28% of variance (R2 = 0.28, F(1, 32) = 12.69, p = 0.001, adj. R2 = 0.26)

### Behavioural and gene expression responses of Astrangia poculata to thermal challenges

Both aposymbiotic and symbiotic colonies reduced their polyp activity in response to food under cold challenge (p < 0.001) and heat challenge (p = 0.012) relative to fragments under control conditions, and this response was most pronounced when temperatures approached their extremes towards the end of the experiment (Figure 1B). A significant interaction between symbiotic state and heat challenge (p = 0.003) was also observed with symbiotic fragments exhibiting less polyp activity than aposymbiotic fragments when temperatures increased. No interaction between symbiotic state and cold challenge was observed (p = 0.083).

Both thermal challenges elicited strong transcriptome-wide changes in gene expression in *A. poculata* (Figure 2A, *Adonis* ptreatment < 0.001). Corals in the cold challenge exhibited greater transcriptome plasticity (Figure 2B, F(1, 22) = 143.35, p < .001, 95% CI [0.77, 1.00]) relative to those under heat challenge. This plasticity corresponded to approximately six times as many DEGs (FDR < 0.05) in cold challenge (6,690 (19.9% of total genes) DEGs, 2,549 (7.6%) upregulated, 4,141 (12%) downregulated) compared to heat challenge (1,011 (3%) DEGs; 552 (1.6%) upregulated, 459 (1.4%) downregulated).

**Figure 2.**
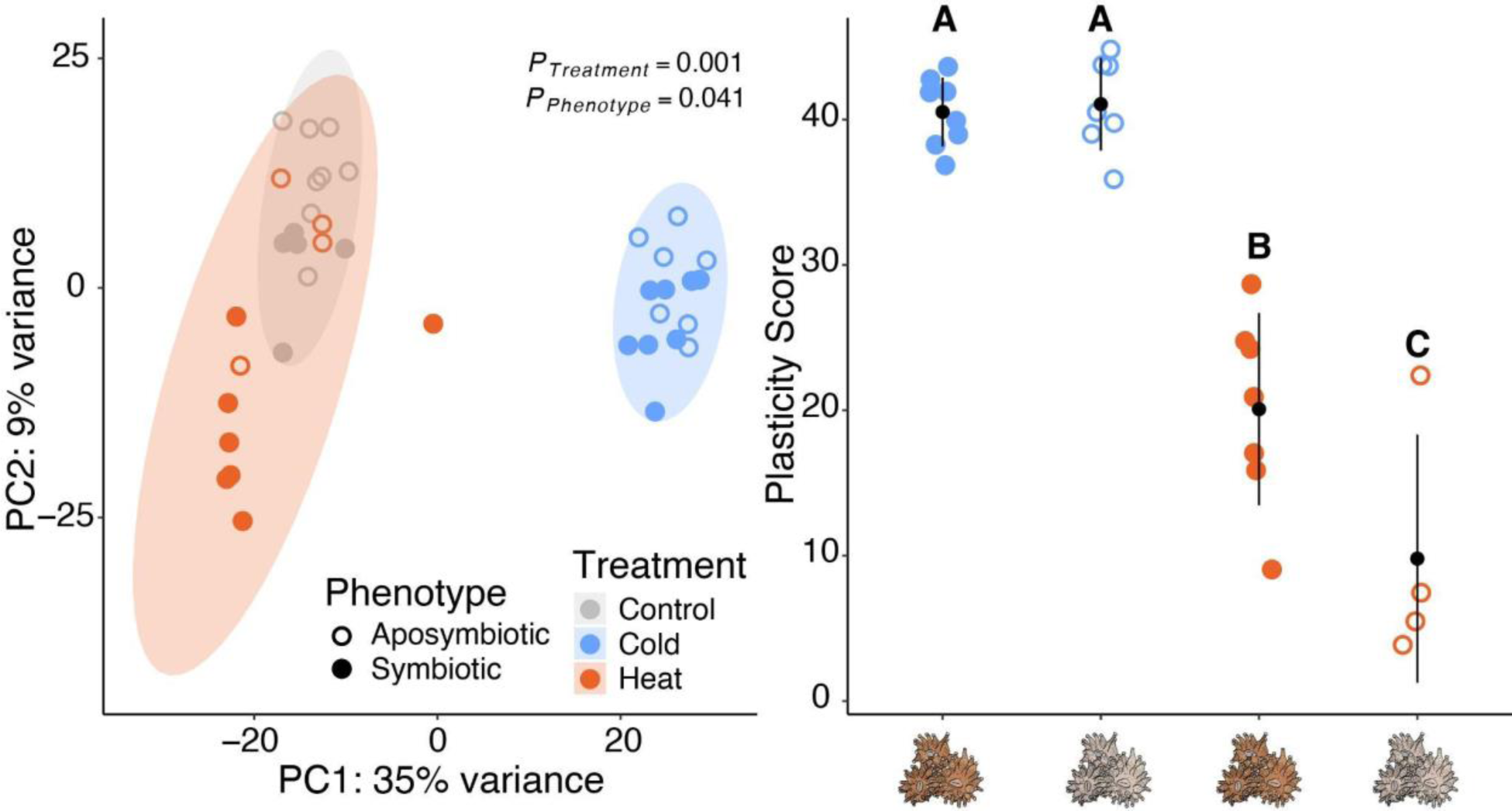
Gene expression responses to temperature challenges across symbiotic states in *Astrangia poculata*. A) Principal component (PC) analysis of global expression of all *A. poculata* vst-normalized genes. Percentages represent the variance explained by each principal component (PC) and shaded areas represent 95% confidence ellipses within treatments. P-values indicate significant main effects of a factor using a permutational multivariate analysis of variance. B) Mean gene expression plasticity of corals in thermal challenge treatments relative to control samples. Plasticity scores represent the distances of each coral fragment in a thermal challenge relative to the average expression of all control fragments across the first two PCs. Symbol and error bars are the modelled means and 95% confidence intervals. Letters depict significant differences in gene expression plasticity across treatments and symbiotic states based on Tukey’s honest significant differences post hoc test.

### Response to heat challenge in Astrangia poculata is mediated by symbiotic state

Gene expression plasticity was significantly higher in symbiotic corals compared to aposymbiotic corals under heat challenge (Tukey’s HSD, p = 0.0203; Figure 2), and these patterns were confirmed by DAPC (Tukey’s HSD, p = 0.0047; Figure S5). These differences in gene expression plasticity were consistent with the number of DEGs (Figure 3), where symbiotic corals had 558 (1.6%) DEGs (351 (1%) upregulated, 207 (0.62%) downregulated) under heat challenge compared to only 172 (0.5%) in heat-challenged aposymbiotic corals (61 (0.18%) upregulated, 111 (0.33%) downregulated).

**Figure 3.**
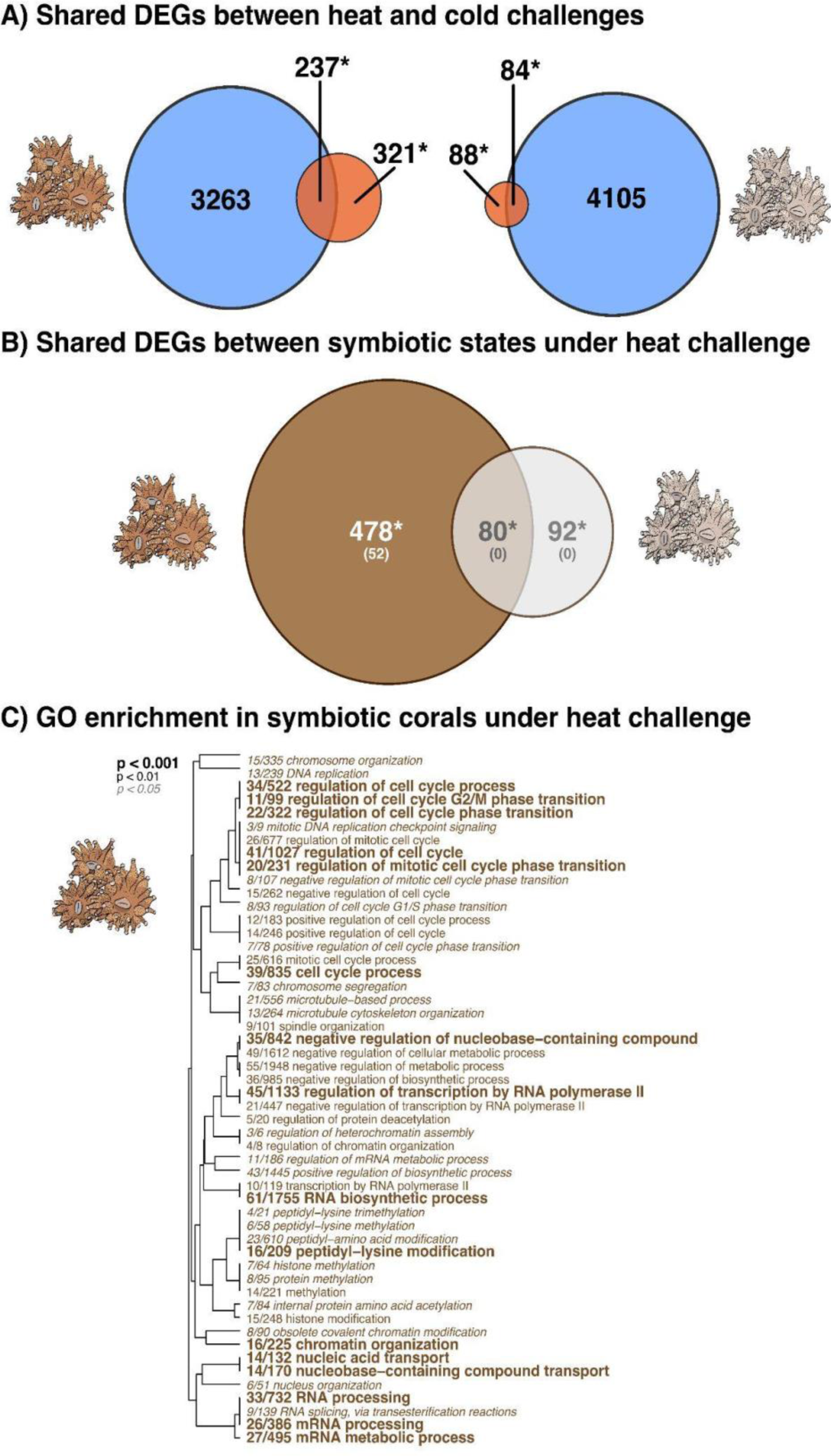
Functional responses of thermal-challenged *Astrangia poculata*. A) Venn diagram of differentially expressed genes (DEGs, FDR < 0.05) from symbiotic (left) and aposymbiotic (right) *A. poculata* fragments under heat challenge (red + asterisk) and cold challenge (blue) relative to control conditions. B) Venn diagram of DEGs from the heat challenge (from asterisks above) between symbiotic (brown) and aposymbiotic (grey) colonies. The top number denotes the number of DEGs and bottom number represents the number of enriched gene ontology (GO) terms. C) GO enrichment results of the “biological processes (BP)” category derived from the list of unique DEGs that responded to heat challenge in the symbiotic corals only. The dendrogram describes the relationship of shared genes between categories, and text size and boldness indicates the significance of each term.

No GO enrichment was observed in the list of DEGs unique to heat-challenged aposymbiotic corals or those genes shared between both symbiotic and symbiotic corals under heat challenge (Figure 3). In contrast, DEGs unique to heat-challenged symbiotic corals were significantly enriched (FDR < 0.05; BP = 52, Figure 3B; CC = 19, Figure S7B; MF = 13, Figure S7C)) with two distinct clusters of related GO terms. The first cluster of terms was related to growth regulation (*e.g.,* GO:0051726: regulation of cell cycle, GO:0022402: cell cycle process, GO:1901990: regulation of mitotic cell cycle phase transition) and the second the formation and metabolism of RNA (*e.g.*, GO:0016071: mRNA metabolic process, GO:0032774: RNA biosynthetic process, GO:0018205: peptidyl-lysine modification, GO:0006357: regulation of transcription by RNA polymerase II, GO:0006325: chromatin organization).

### Characterising the environmental stress response in Astrangia poculata

To explore differential regulation of genes associated with oxidative stress, delta ranks of 13 children terms belonging to the parent GO term *oxidative stress* (GO:0006979) were explored. Delta-ranks of these children terms elicited similar responses under both heat and cold challenge (F(1, 48) = 0.38, p = 0.538), and this similarity was maintained across symbiotic states (Figure 4; F(1, 48) = 1.07, p = 0.308), suggesting no change in the oxidative stress response across thermal challenges or symbiotic states.

**Figure 4.**
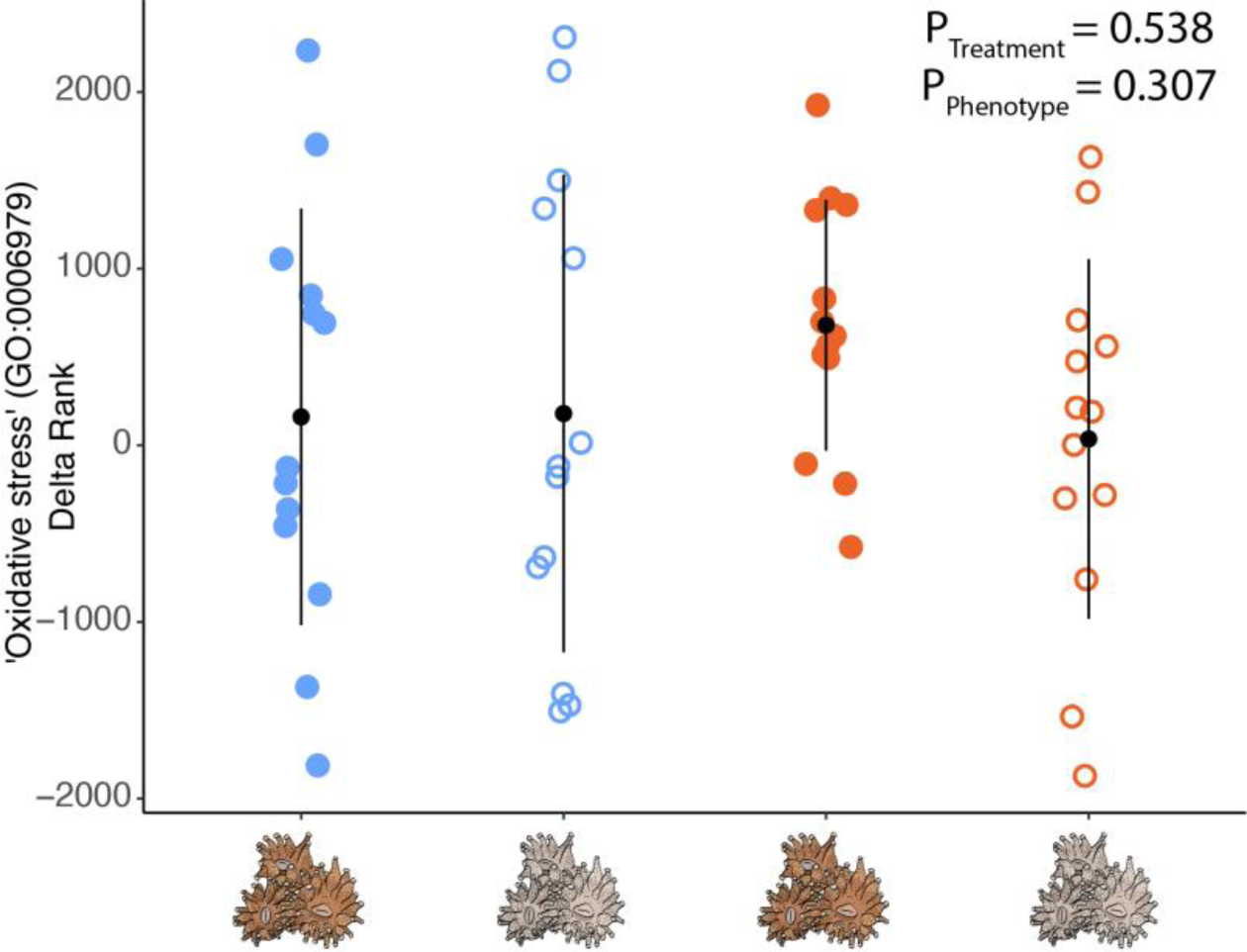
The oxidative stress response does not change between symbiotic states. Each dot represents the delta rank of one of the 13 children gene ontology (GO) terms under the oxidative stress parent term (GO:0006979) across thermal challenges within each symbiotic state. Black dots represent mean delta rank values and error bars denote standard deviation.

To further explore the ESR beyond oxidative stress, delta ranks from Mann-Whitney U GO enrichment tests for each symbiotic state and thermal challenge were contrasted with the delta ranks of the Type A stress response studies from Dixon *et al.* (2020). While not a formal statistic, a generally positive relationship suggests functional similarities between our data and that of the Type A stress studies in Dixon *et al.* (2020). There was a positive relationship for both symbiotic states under cold challenge in the ‘biological processes’ GO category (Figure 5B,D), suggesting a “Type A” stress response. This positive slope was consistent in the ‘molecular function’ and ‘cellular component’ categories as well (Figure S6 B-C). In contrast, heat challenge elicited more dissimilar GO functionality as there was an opposing slopes for aposymbiotic and symbiotic *A. poculata*, with aposymbiotic corals exhibiting a positive slope (Type A) across all GO categories while symbiotic corals showcased a “Type B” stress response as indicated by the negative slope (Figure 5C [‘biological processes’ GO category], Figure S7 B-C [‘molecular function’ and ‘cellular component’ GO categories]).

**Figure 5.**
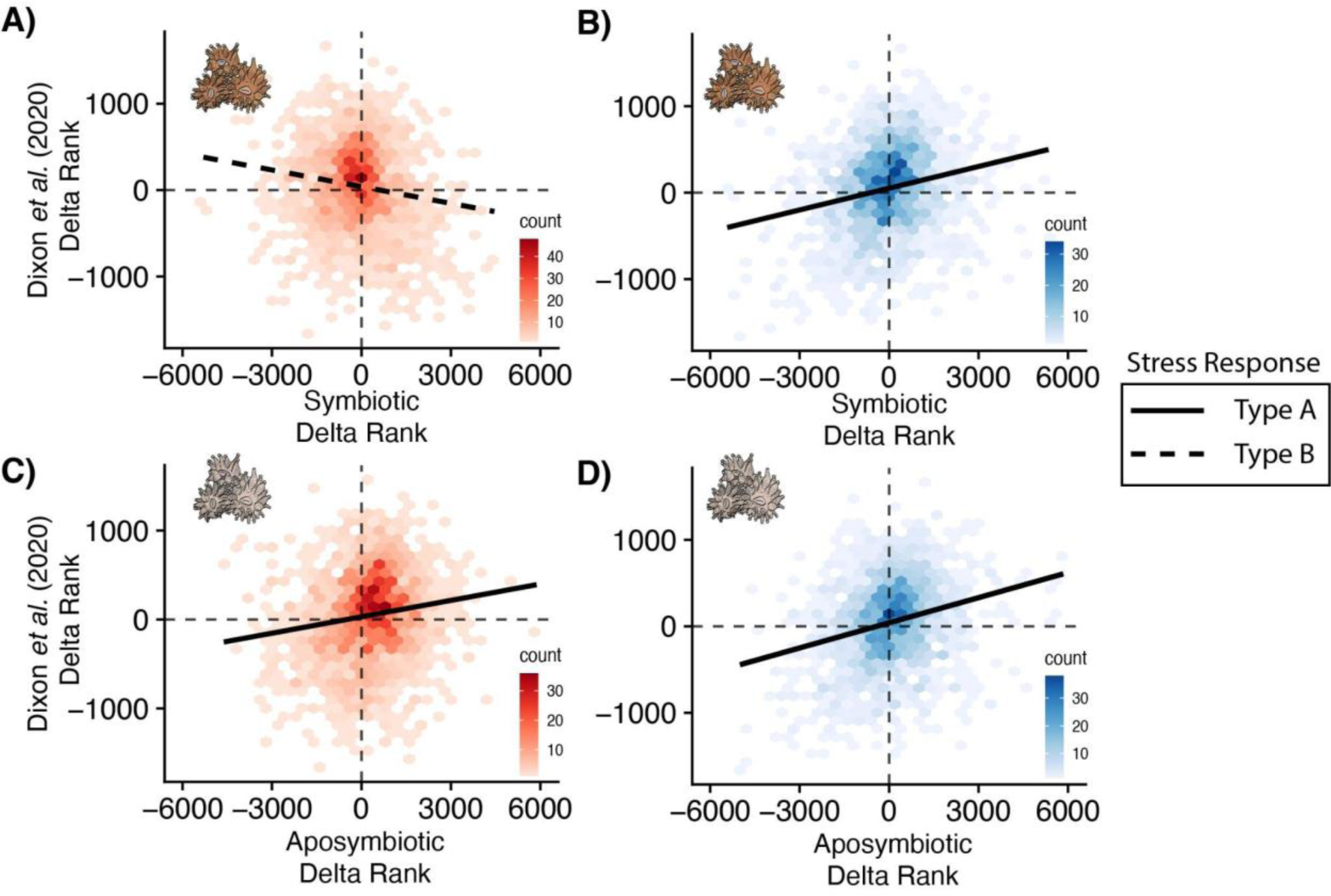
Environmental stress responses (ESR) of *Astrangia poculata*. Comparison of functional similarities in gene ontology (GO) delta ranks from studies that exhibited a Type A stress response (Dixon et al., 2020) compared with those from the heat (A, C) and cold challenge delta ranks (B, D) from aposymbiotic (C,D) and symbiotic states (A, B). Functional GO enrichment are shown for the biological processes (BP) and functional similarities between the two data sets would be characterised by a positive slope indicates that corals in a thermal challenge are eliciting a functionally similar response to that of tropical corals that have a severe ESR (Type ‘A’) whereas a negative slope suggests that the responses are more dissimilar therefore more similar to tropical corals exhibiting a moderate ESR (Type ‘B’).

## Discussion

Here, we conducted thermal challenge experiments on aposymbiotic and symbiotic fragments of the facultative symbiotic coral *Astrangia poculata* with the broad goal of testing the influence of photosymbiosis on thermal stress responses. We predicted that under the oxidative bleaching hypothesis [23] (Table 1), there would be an elevated environmental stress response (ESR; Type A response) due to the accumulation of photobiont and host derived reactive oxygen species (ROS) when thermally challenged. We found that photosymbiosis differentially modulates the coral’s response to heat challenge; however, this response was not consistent with the expectations of the oxidative bleaching hypothesis. Instead, our results highlight that coral gene expression patterns showcase enrichment for pathways involved in the regulation of symbiont growth *via* host cell cycle mechanisms under heat challenge. In addition, comparative analyses with a previous gene expression meta-analysis in tropical corals suggest that photosymbiosis was associated with a less strong ESR (Type B response) under heat challenge in this facultative coral system. In contrast, under cold challenge, *A. poculata* exhibited stronger responses compared to heat challenge and photosymbiosis did not impact these gene expression responses.

*Astrangia poculata* experiences wide seasonal variation in temperature, including winter temperatures that are colder than the cold challenge applied here (Figure 1C). Despite this, cold challenge elicited strong behavioural and transcriptomic responses in both aposymbiotic and symbiotic colonies (Figure 2; Figure S5). While these results align with our previous findings in aposymbiotic *A. poculata* [22], they also showcase that photosymbiosis failed to modulate this response. This pattern contrasts with a study in *A. poculata* that found that photosymbiosis mitigated host physiological responses under cold temperatures (8°C), with symbiotic colonies healing more quickly than aposymbiotic colonies [34]. It is possible that the colder temperatures (4°C) used in our study relative to [32] elicited a stronger cold response, and this may have swamped out the effects of photosymbiosis. Nevertheless, the consistently strong gene expression plasticity under cold challenge is relevant given that cold water from arctic currents constrain *A. poculata* from expanding beyond Cape Cod [61]. Extreme cold weather therefore remains a salient stressor for *A. poculata* under climate change as Arctic warming influences upper-level atmospheric activity, which fuels severe winters [62]. These cold winters will therefore likely continue to constrain range expansion north in this species even as climate change progresses.

While photosymbiosis had little impact on host gene expression under cold challenge, previous studies have highlighted how photobiont biology is impacted by colder temperatures. The photobiont of *A. poculata* (*Breviolum psygmophilum*) exhibited reduced photosynthetic efficiency (Fv/Fm) as temperatures decreased, both *in hospite* [34] and *ex hospite* [63]. While the *ex hospite* work initially showed a decrease in Fv/Fm, *B. psygmophilum* showed resilience and Fv/Fm quickly recovered once temperatures returned to baseline [63]. While corals were not offered a recovery period in our study, this pattern in *A. poculata’s* photobiont suggests that hosts may also demonstrate similar resilience if they are co-adapted to their environments, which has been shown in other obligate symbiotic corals [64]. It has also been shown that *in hospite A. poculata* photobionts experience large seasonal variation in cell densities with lower photobiont densities observed in winter months [39]; however, this reduction in cell density can take several months of cold temperatures to manifest [65]. While cold challenge experiments are less common in coral reef studies given the imminent threat of ocean warming, previous work on photobiont physiology has showcased mixed results in tropical and subtropical coral species. In contrast to Sharp *et al.* [39], a 10 week cold exposure (23°C; control 26°C) in *Acropora millepora* increased photobiont densities [12], while a more severe cold challenge in tropical and subtropical *Porites lutea* populations (temperatures lowered from 26°C-12°C over 18 days) reduced photobiont densities and photosynthetic efficiencies [66]. Clearly, the rate of cooling and severity of the temperature achieved likely dictate the magnitude and severity of photobiont responses and highlights the importance of these aspects in experimental studies [67]. It is also critical to point out that the rate of temperature change used here does not represent an ecologically relevant change for *A. poculata* given that these corals experience large differences in seasonal temperatures that occur over longer timescales.

Gene expression plasticity in response to cold challenge across both symbiotic states was notably large (Figure 2), although no difference between symbiotic states was observed. We speculate that our sampling of *A. poculata* from their northern range edge may explain this plasticity. Plasticity can allow populations to establish in marginal habitats as they expand their ranges, so sampling along these range edges could be biased towards individuals with high plastic [68,69]. Equally plausible, high gene expression plasticity may be explained by the climate variability hypothesis (CVH), which proposes that organisms experiencing high seasonal variability will exhibit higher plasticity facilitating acclimation across broad thermal regimes [70]. Unfortunately, our experiment is unable to discern the mechanisms underlying this high gene expression plasticity under cold challenge. Nevertheless, this pattern is consistent with previous work on high latitude, marginal populations of *Porites lutea,* which were found to not only exhibit higher cold tolerance, but also stronger transcriptomic responses when subjected to cold challenge relative to those sampled from the core tropical range [66]. Future comparative work on multiple populations of *A. poculata* is warranted.

Relative to cold challenge, we observed muted gene expression responses to heat challenge despite exceeding temperatures experienced at the collection site (Figure 1). A large plastic response to cold, but not heat challenge is seemingly at odds with both the marginal environment hypothesis and the CVH, as both would predict highly plastic responses to both temperature treatments. However, it is possible that short term cold challenges are more stressful for *A. poculata* than short term heat challenges. For example, experimental work on the tropical coral *Acropora yongei* found that short term cold stress was more detrimental than short term heat stress, but longer-term elevated temperatures were ultimately more harmful than longer-term colder temperatures [71]. Alternatively, the rate of cooling may have a stronger influence than the rate of warming on *A. poculata.* Given that this population of this species is known to enter a hibernation-like state called quiescence after long seasonal decreases in temperature [72], perhaps more gradual temperature decreases would induce acclimatory responses that lead to quiescence rather than the strong ESR observed here. In addition, these colonies were acclimated to ambient room temperatures which is relatively warmer than the average temperatures experienced from their collection site. This acclimation may impact their response to the heat challenge provided here, and future experimental work comparing responses in winter-*versus* summer-acclimated colonies is needed. Lastly, we posit that this muted response to heat challenge may be further evidence that Woods Hole, MA represents a marginal habitat for *A. poculata* and that these corals are adapted to more subtropical locations (*e.g.* Virginia). Indeed, high gene flow between southern (*i.e.,* North Carolina) and northern (*i.e.,* Massachusetts) populations has been documented [73], highlighting a need for future reciprocal transplant experiments between these locations to test this hypothesis. In response to heat challenge, symbiotic *A. poculata* exhibited higher gene expression plasticity and approximately three times as many DEGs as aposymbiotic *A. poculata* (Figure 3). GO enrichment analyses of these DEGs unique to symbiotic colonies were used to functionally interpret the pathways that were enriched under heat challenge. This analysis largely highlighted genes involved in cell cycle processes (Figure 3). This interaction between photobionts and hosts leading to changes in cell cycle processes is relevant as the division of host and photobiont cells are often synchronized [74], preventing photobiont overpopulation shifting the mutualism to parasitism [75]. Furthermore, one mechanism that hosts use to modulate photobiont growth is by maintaining the space available for photobiont cells to grow into (for review, see [78]). Similar GO enrichment between symbiotic and aposymbiotic tissues have been observed in the facultatively symbiotic coral *Oculina arbuscula* under baseline conditions [42], suggesting that controlling symbiont cell densities is a critical aspect of symbiosis maintenance. Interestingly, research has shown that cell densities can increase under short term thermal challenge, which can lead to high symbiont loads that may increase ROS production within hosts [77]. This contrasts with recent work suggesting that corals can farm and digest excess symbiont cells [79], which would simultaneously increase nutrient availability to the host and control cell densities. Overall, we posit that under heat challenge, *A. poculata* exhibits enrichment of cell cycle processes, which are necessary to control symbiont cell densities under warmer temperatures. Whether or not these corals control cell densities via symbiont cell digestion remains to be tested.

At first, we hypothesised that the observed elevated gene expression plasticity under heat challenge in symbiotic corals would be accompanied by a greater ESR associated with accumulation of photobiont derived ROS (Table 1). Indeed, experimental evidence supports photobionts imposing oxidative stress on cnidarian hosts [23,80,81]. For example, ROS leakage from freshly isolated photobionts in a heat stress experiment can increase by up to 45%, correlating with increases in coral host oxidative damage [82]. However, further inspection of genes belonging to GO terms nested within “oxidative stress” demonstrated no differences in the enrichment of these terms between aposymbiotic and symbiotic colonies under either thermal challenge (Figure 4). Therefore, this pattern does not support the hypothesis of increased ROS produced from algal photobionts in symbiotic corals under heat challenge. This pattern could be due to constitutively higher antioxidant and stress mitigation mechanisms under photosymbiosis, perhaps mediated through energetic increases associated with symbiont digestion [79]. For example, melanin and catalase levels, which play roles in mitigating stress, are higher in symbiotic *A. poculata* than aposymbiotic colonies [35]. While it is possible that by sampling for gene expression at the end of the challenge we were unable to capture a signal of increased oxidative stress that may have been present earlier in the experiment, it is also equally plausible that this pattern supports the growing evidence that ROS accumulation from photoinhibition may not be the primary mechanism of coral bleaching [83]. For example, ROS accumulation in *Aiptasia* after a heat shock was determined to be host derived and generated prior to photobiont photoinhibition [84]. Furthermore, when exogenous antioxidants are treated to *Aiptasia* under elevated stress, the anemones still bleached despite having no changes in relative ROS [85]. Furthermore, coral bleaching can occur under complete darkness, suggesting that this process can be independent of photosynthesis [86]. Taken together, these results have reshaped our understanding of bleaching, shifting from the notion of photo-induced ROS accumulation and toward the recognition of nutritional mechanisms potentially altering responses to thermal stress [28]. However, it is important to acknowledge that temperate corals may exhibit inherently different physiology than tropical corals as they are less reliant on photosynthetically derived sources of energy.

Lastly, to further explore the ESR beyond oxidative stress, we compared our findings with a meta-analysis of coral gene expression responses to stress [21]. This comparative work facilitated broadly assessing the type of ESR responses elicited by the thermal challenges between symbiotic states (Figure 5). Interestingly, we observed divergent ESRs to heat challenge between symbiotic states, with expression patterns of aposymbiotic corals consistent with a ‘Type A’ response (positive slope) and symbiotic corals a ‘Type B’ response (negative slope). Type B responses are typically observed under moderate stress [21], suggesting that symbiotic *A. poculata* exhibited a less severe ESR than aposymbiotic corals under heat challenge. This result contrasts previous work demonstrating that aposymbiotic *A. poculata* had Type B response to heat challenges [22], and higher thermal optima compared to symbiotic corals [87]. However, [87] did not capture the full thermal performance curve, and molecular stress signatures are expected to occur outside of the thermal optimum [88]. Future work exploring whether thermal challenges beyond a population’s critical thermal maximum shift this ESR under heat challenge is warranted. Ultimately, the mechanisms underlying how photosymbiosis reduces the ESR under heat challenge remain to be determined. Photobionts may lessen stress by protecting the coral from the compounding effects of light stress. As photobionts themselves are pigmented, they block or absorb light that would otherwise be scattered and amplified by the coral skeleton [89,90], potentially limiting the additive effects of temperature and light stress. Alternatively, photobionts provide carbon sugars to the host, so this additional energy input may mitigate ESRs in symbiotic corals under heat challenge [30]. Alternatively, hosts may farm and consume these algae for excess nutrition [79]. Future work aiming to disentangle how photosymbiosis mitigates the ESR in symbiotic *A. poculata* would benefit from a deeper understanding of 1) *A. poculata*’s thermal maximum, 2) how the rate of temperature increases impacts physiology, 3) nutrient exchange mechanisms in this facultative symbiosis (but see [37]), and 4) how changes in symbiont density shift these responses. Overall, these findings showcase the potential complexities of photosymbiosis in this system and fail to implicate the symbiont in amplifying the coral ESR under heat.

## Acknowledgements

This work stemmed from the collaborative efforts made possible by Boston University’s Marine Program (BUMP). The authors extend deep appreciation to Justin Scace, Julia Mendez and Nicola Kriefall who facilitated the experiential learning course as well as students including Kalli Richmond in the BUMP program who provided husbandry and experimental assistance. We also thank Kathryn Stankiewicz and Iliana Baums for providing early access to the *Astrangia poculata* genome for mapping our data. We acknowledge Boston University’s Shared Computing Cluster (SCC) for computational resources associated with these analyses. Members of the Davies lab are also acknowledged for their thoughtful feedback and perspectives from Pete Buston, Randi Rotjan, Sean Mullen and Rachel Wright throughout the analysis, interpretation, and editing process. The authors extend appreciation to the annual Temperate Coral Research Conferences hosted by Roger Williams University, Boston University, and Southern Connecticut State University for fostering creative conversations and collaborations leading to this work.

## Data Availability Statement

Raw sequences are made available from the NCBI SRA under accession PRJNA1013245. Full reproducible data and code as well as all intermediate files are available at https://github.com/wuitchik

