## Supplemental Figures for "Photosymbiosis reduces the environmental stress response under a heat challenge in a facultatively symbiotic coral"

**Supplemental Information**

Table S1 | Sample size post filtering and outlier removal used for gene expression analyses

| | Heat Challenge | Control | Cold Challenge |
|---|---|---|---|
| Symbiotic | 7 | 5 | 8 |
| Aposymbiotic | 4 | 8 | 7 |

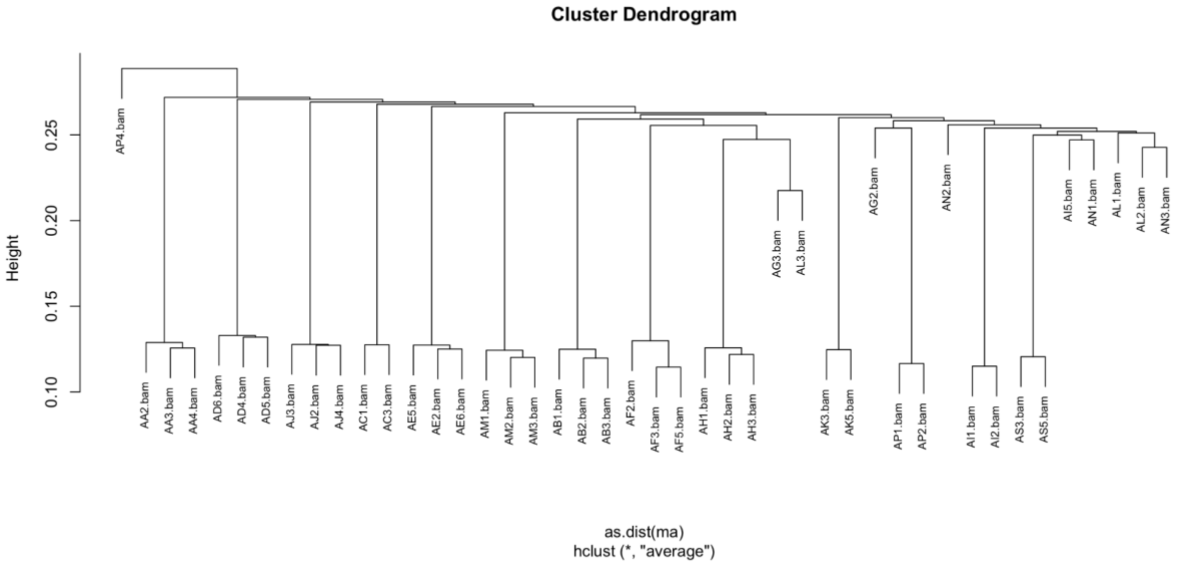


Figure S1| Cluster dendrogram of *Astrangia poculata* to identify clonal individuals from SNPs. Sample AP4 was removed from analyses due to its high divergence between all other samples.

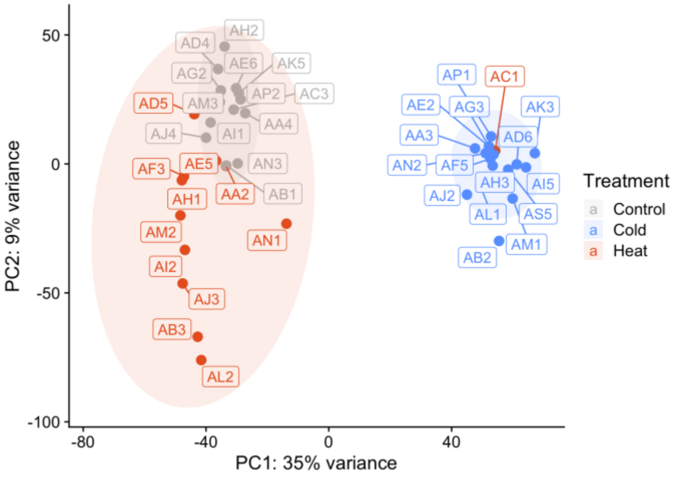


Figure S2. Gene expression responses to temperature challenge treatments prior to sample “AC1” removal. Principal component analysis of overall expression of all *A. poculata* vst-normalized genes. Percentages represent the variance explained by each axis and shaded areas represent 95% confidence ellipses within treatments. Sample AC1 was removed due to its high divergence in gene expression profile.

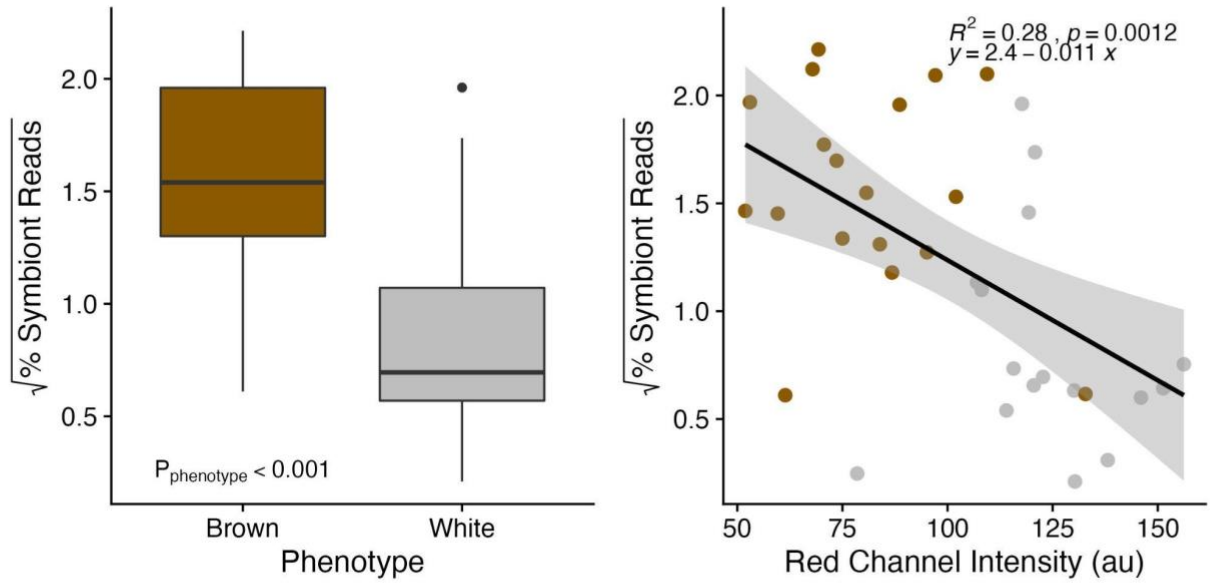


Figure S3. Association of reads mapped to photobiont and phenotype. Left) Subjective phenotype (Brown or White) Right) Calculated red channel intensity from image analysis.

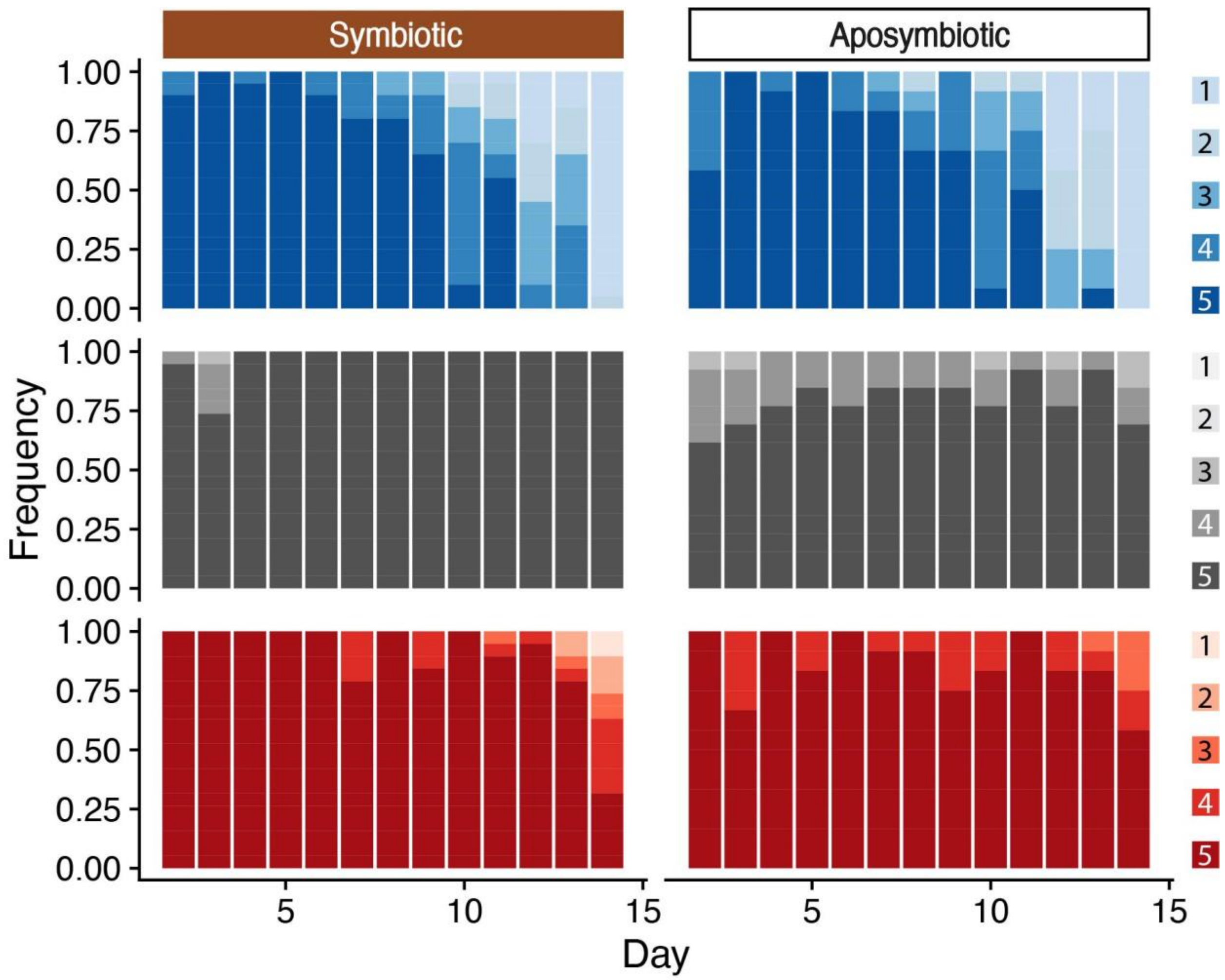


**Figure S4.** Behavioural response to food stimuli. Polyp activity was measured as the proportion of polyps that were extended per coral fragment (approximately, 1 = 0%, 2 = 25%, 3 = 50%, 4 = 75%, 5 = 100%) across the 15 day experiment. Cold challenge (Blues; top), control (Greys; middle) and heat challenge (Reds; bottom) with respective legend showing behavioural scores.

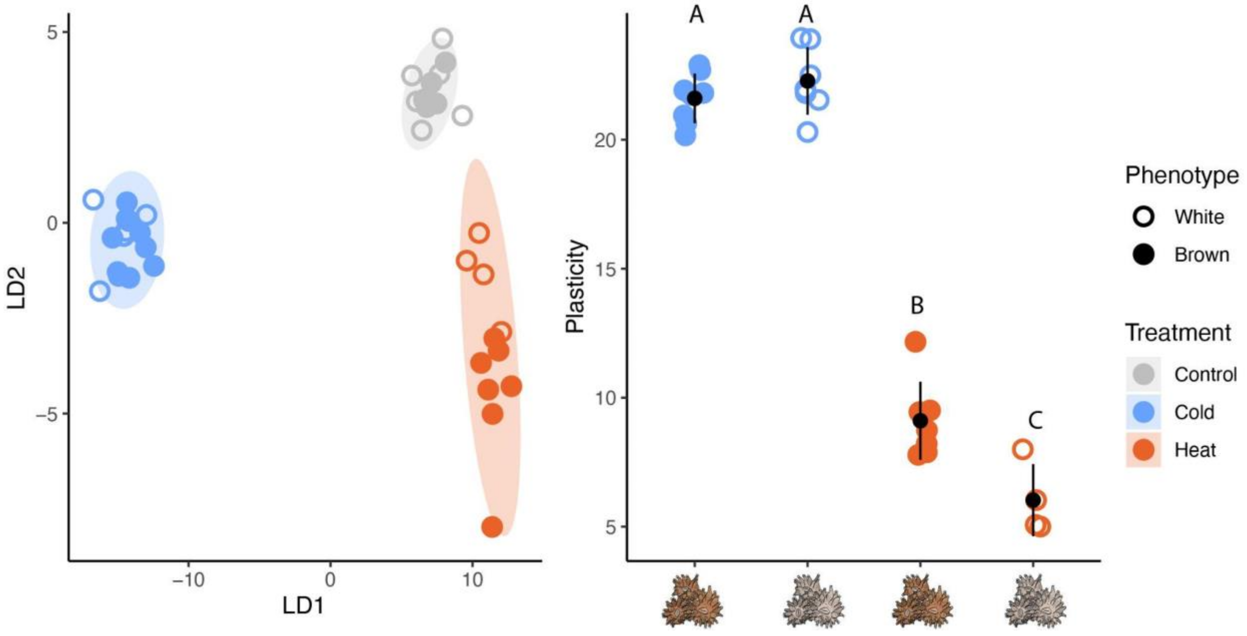


Figure S5. A) Discriminant function analysis of overall expression of all *A. poculata* rlogged-normalized genes. Shaded areas represent 95% confidence ellipses within treatments. B) Mean gene expression plasticity of corals in thermal challenge treatments relative to control samples. Plasticity scores represent the first two principal component distances of each coral fragment in a thermal challenge treatment relative to the average expression of all control fragments. Symbol and error bars are the modeled means and 95% confidence interval. Letters depict significant differences in gene expression plasticity across treatments and symbiotic states based on Tukey's honest significant differences post hoc test.

Table S2 | Summary of behavioural ordinal logistic regression treating genotype and system as random effects.

| | **Estimate** | **Std. Error** | **z value** | **Pr(>\|z\|)** |
|---|---|---|---|---|
| Cold Challenge | 4.886 | 0.835 | 5.854 | **<0.001** |
| Heat Challenge | 1.960 | 0.783 | 2.503 | **0.012** |
| Symbiotic State | -1.035 | 1.228 | -0.844 | 0.399 |
| Experimental day Day | 0.059 | 0.053 | 1.116 | 0.264 |

| | | | | |
|---|---|---|---|---|
| Cold Challenge * Symbiotic State | 2.653 | 1.531 | 1.733 | 0.083 |
| Heat Challenge * Symbiotic State | 5.460 | 1.815 | 3.009 | **0.003** |
| Cold Challenge * Experimental Day | -0.771 | 0.089 | -8.624 | **<0.001** |
| Heat Challenge * Experimental Day | -0.188 | 0.085 | -2.212 | **0.027** |
| Symbiotic State * Day | 0.782 | 0.354 | 2.208 | **0.027** |
| Cold Challenge * Symbiotic state * Experimental Day | -0.904 | 0.366 | -2.470 | **0.014** |
| Heat Challenge * Symbiotic State * Experimental Day | -1.179 | 0.375 | -3.146 | **0.002** |

**Table S3.** Read counts and sample notes at various steps of the mapping process.

| Sample | Genotype | Treatment | Phenotype | Raw total reads | Trimmed total reads | Mapped reads to host | Percent mapped to host | Photobiont counts | Relative Photobiont Counts | Notes |
|---|---|---|---|---|---|---|---|---|---|---|
| AA2 | A | Heat | White | 5057078 | 1389360 | 970628 | 70% | 5929 | 0.60% | |
| AA3 | A | Cold | White | 3740666 | 876102 | 574992 | 66% | 2309 | 0.40% | |
| AA4 | A | Control | White | 2780124 | 837632 | 595368 | 71% | 11632 | 2.00% | |
| AB1 | B | Control | Brown | 5130310 | 1399027 | 1010691 | 72% | 30109 | 3.00% | |
| AB2 | B | Cold | Brown | 5607990 | 1568156 | 585488 | 37% | 23201 | 4.00% | |
| AB3 | B | Heat | Brown | 810976 | 301017 | 226910 | 75% | 10972 | 4.80% | |
| AC1 | C | Heat | White | 6186284 | 1808120 | 1245649 | 69% | 3396 | 0.30% | Removed based on PCA |
| AC3 | C | Control | White | 8772244 | 2254587 | 1535196 | 68% | 6770 | 0.40% | |
| AD4 | D | Control | White | 5621719 | 1178831 | 834049 | 71% | 1264 | 0.20% | |
| AD5 | D | Heat | White | 2609448 | 702086 | 269066 | 38% | 7010 | 2.60% | |

| | | | | | | | | | | |
|---|---|---|---|---|---|---|---|---|---|---|
| AD6 | D | Cold | White | 6092666 | 1545354 | 1170859 | 76% | 6111 | 0.50% | |
| AE2 | E | Cold | White | 3056318 | 1042104 | 777881 | 75% | 1042 | 0.10% | |
| AE5 | E | Heat | White | 4528236 | 1193039 | 546344 | 46% | 4775 | 0.90% | |
| AE6 | E | Control | White | 5024737 | 1294243 | 762508 | 59% | 3929 | 0.50% | |
| AF2 | F | Control | Brown | 8447 | 4222 | 3509 | 83% | 46 | 1.30% | Removed due to low counts and because of identified outlier on arrayQualitymetrics |
| AF3 | F | Heat | Brown | 3844256 | 949292 | 546121 | 58% | 2442 | 0.40% | |
| AF5 | F | Cold | Brown | 5017655 | 1394088 | 976195 | 70% | 24009 | 2.50% | |
| AG1 | G | Heat | White | 51 | 28 | 21 | 75% | 0 | 0.00% | Removed due to low counts |
| AG2 | G | Control | White | 5632433 | 1586345 | 1141961 | 72% | 41035 | 3.60% | |
| AG3 | G | Cold | White | 6128832 | 1645995 | 1257027 | 76% | 15370 | 1.20% | |
| AH1 | H | Heat | White | 4562147 | 940299 | 665266 | 71% | 3100 | 0.50% | |
| AH2 | H | Control | White | 2653474 | 454304 | 340981 | 75% | 1883 | 0.60% | |
| AH3 | H | Cold | White | 2471788 | 790449 | 609247 | 77% | 8044 | 1.30% | |
| AI1 | I | Control | Brown | 4720194 | 1334265 | 1009054 | 76% | 11998 | 1.20% | |
| AI2 | I | Heat | Brown | 1101842 | 431060 | 318267 | 74% | 12738 | 4.00% | |
| AI5 | I | Cold | Brown | 1655390 | 630365 | 487477 | 77% | 8760 | 1.80% | |
| AJ2 | J | Cold | Brown | 2983572 | 955180 | 689278 | 72% | 9467 | 1.40% | |
| AJ3 | J | Heat | Brown | 2565084 | 775431 | 557771 | 72% | 23536 | 4.20% | |
| AJ4 | J | Control | Brown | 4827011 | 940906 | 613723 | 65% | 5264 | 0.90% | |
| AK2 | K | Heat | White | 107 | 70 | 38 | 54% | 1 | 2.60% | Removed due to low counts |
| AK3 | K | Cold | White | 3009774 | 690136 | 511510 | 74% | 1143 | 0.20% | |
| AK5 | K | Control | White | 3280922 | 1012242 | 768666 | 76% | 4163 | 0.50% | |

| | | | | | | | | | | |
|---|---|---|---|---|---|---|---|---|---|---|
| AL1 | L | Cold | Brown | 3298436 | 979125 | 665532 | 68% | 17685 | 2.70% | |
| AL2 | L | Heat | Brown | 4785674 | 569249 | 434299 | 76% | 10598 | 2.40% | |
| AL3 | L | Control | Brown | 27109 | 10644 | 8522 | 80% | 240 | 2.80% | Removed due to low counts |
| AM1 | M | Cold | Brown | 2797047 | 609399 | 374660 | 61% | 6682 | 1.80% | |
| AM2 | M | Heat | Brown | 3510007 | 1041855 | 748704 | 72% | 10108 | 1.40% | |
| AM3 | M | Control | Brown | 5028286 | 1193612 | 866928 | 73% | 3290 | 0.40% | |
| AN1 | N | Heat | Brown | 3444755 | 1130026 | 851746 | 75% | 19778 | 2.30% | |
| AN2 | N | Cold | Brown | 4278155 | 1368617 | 1020334 | 75% | 40635 | 4.00% | |
| AN3 | N | Control | Brown | 3587592 | 847269 | 555903 | 66% | 16331 | 2.90% | |
| AP1 | P | Cold | White | 4284600 | 1339960 | 882049 | 66% | 2883 | 0.30% | |
| AP2 | P | Control | White | 1997983 | 626374 | 482611 | 77% | 1937 | 0.40% | |
| AP4 | P | Heat | White | 2456638 | 805254 | 569111 | 71% | 4428 | 0.80% | Removed due to hclust outlier |
| AS1 | S | Control | Brown | 37 | 24 | 16 | 67% | 0 | 0.00% | Removed due to low counts |
| AS3 | S | Heat | Brown | 4066912 | 417483 | 291188 | 70% | 29108 | 10.00% | Identified as outlier in arrayQuality Metrics |
| AS5 | S | Cold | Brown | 2041873 | 668257 | 475513 | 71% | 9555 | 2.00% | |
| | | | average | 3512487 | 926202 | 634657 | 69% | 9887 | 1.80% | |
| | | | min | 37 | 24 | 16 | 37% | 0 | 0.00% | |
| | | | max | 8772244 | 2254587 | 1535196 | 83% | 41035 | 10.00% | |

Table S4. Summary of behavioural ordinal logistic regression treating genotype and system as random effects.

| | Estimate | Std. Error | z value | Pr(>\|z\|) |
|---|---|---|---|---|
| Cold Challenge | 4.886 | 0.835 | 5.854 | **<0.001** |
| Heat Challenge | 1.960 | 0.783 | 2.503 | **0.012** |
| Symbiotic State | -1.035 | 1.228 | -0.844 | 0.399 |
| Experimental day Day | 0.059 | 0.053 | 1.116 | 0.264 |
| Cold Challenge * Symbiotic State | 2.653 | 1.531 | 1.733 | 0.083 |
| Heat Challenge * Symbiotic State | 5.460 | 1.815 | 3.009 | **0.003** |
| Cold Challenge * Experimental Day | -0.771 | 0.089 | -8.624 | **<0.001** |
| Heat Challenge * Experimental Day | -0.188 | 0.085 | -2.212 | **0.027** |
| Symbiotic State * Day | 0.782 | 0.354 | 2.208 | **0.027** |
| Cold Challenge * Symbiotic state * Experimental Day | -0.904 | 0.366 | -2.470 | **0.014** |
| Heat Challenge * Symbiotic State * Experimental Day | -1.179 | 0.375 | -3.146 | **0.002** |

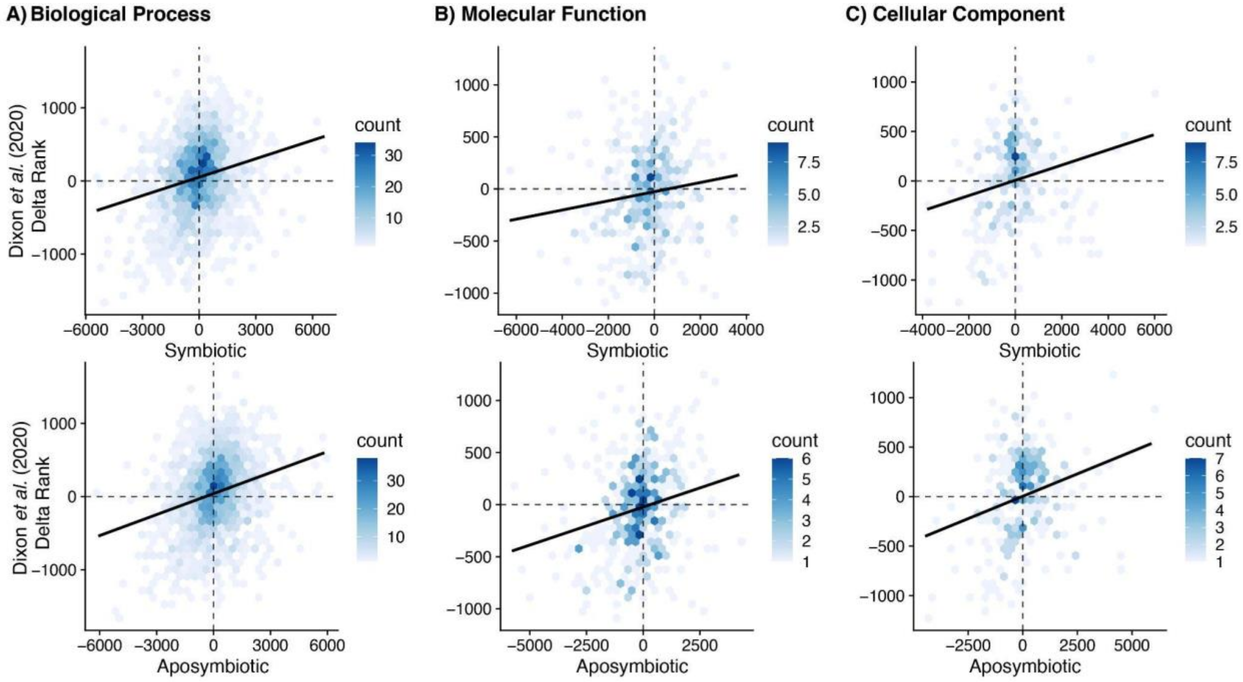


**Figure S6 |** Comparison of gene ontology (GO) delta ranks from environmental stress response studies featured in (G. Dixon et al., 2020) with cold challenge. A positive slope indicates that a coral is showing signatures of severe environmental stress response (ESR) whereas a negative slope is more characteristic of moderate ESR.

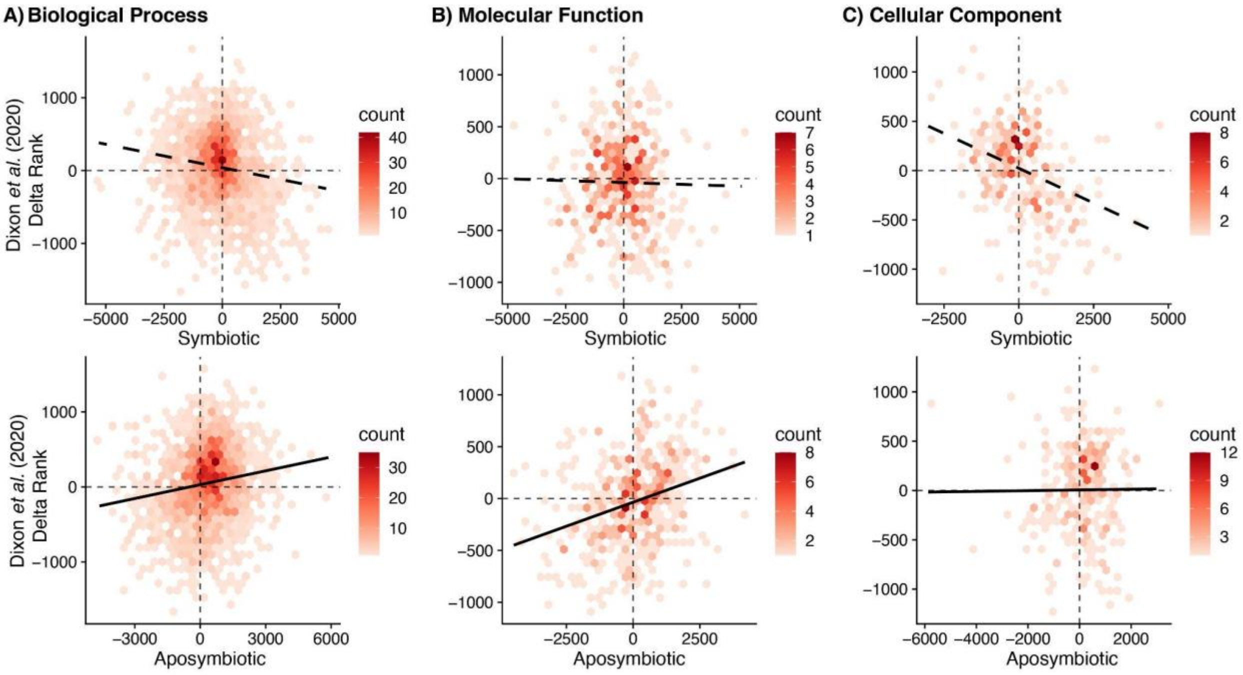

**Figure S7 |** Comparison of gene ontology (GO) delta ranks from environmental stress response studies featured in (G. Dixon et al., 2020) with heat challenge. A positive slope indicates that a coral is showing signatures of severe environmental stress response (ESR) whereas a negative slope is more characteristic of moderate ESR.

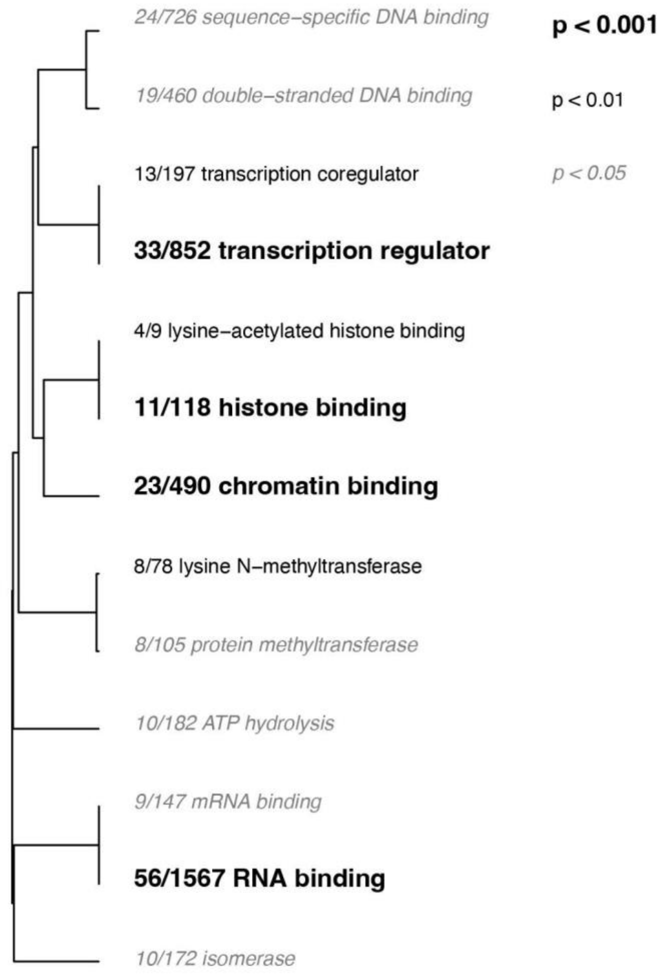

Figure S8 | GO enrichment results of the "molecular functions (MF)" category derived from the list of unique DEGs that responded to heat challenge in the symbiotic phenotype corals only. The dendrogram describes the relationship of shared genes between categories, and text size and boldness indicates the significance of each term.

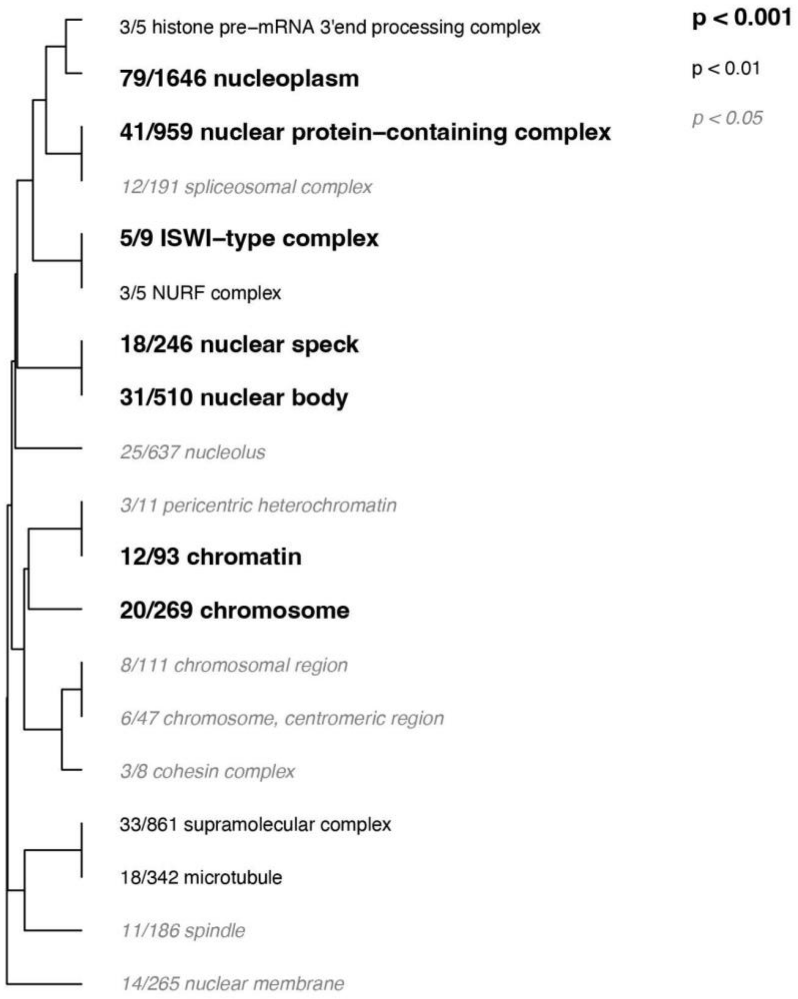

Figure S9 | GO enrichment results of the “cellular components (CC)” category derived from the list of unique DEGs that responded to heat challenge in the symbiotic phenotype corals only. The dendrogram describes the relationship of shared genes between categories, and text size and boldness indicate the significance of each term.
